# Predicting neuronal firing from calcium imaging using a control theoretic approach

**DOI:** 10.1101/2024.11.04.621814

**Authors:** Nicholas A. Rondoni, Fan Lu, Dan Turner-Evans, Marcella Gomez

## Abstract

Two-photon calcium imaging has become a powerful tool to explore the functions of neurons and the connectivity of their circuitry. Frequently, fluorescent calcium indicators are taken as a direct measure of neuronal activity. These indicators, however, are slow relative to behavior, obscuring functional relationships between an animal’s movements and the true neuronal activity. As a consequence, the firing rate of a neuron is a more meaningful metric. Converting calcium imaging data to the firing of a neuron is nontrivial. State of the art methods depend largely on neural networks or non-mechanistic processes, which may yield acceptable correlations between calcium dynamics and the frequency at which a neuron fires, but do not illuminate the underlying chemical exchanges within the neuron or require significant data to be trained on. Leveraging modeling frameworks from chemical reaction networks (CRN) coupled with modern control architectures, a new algorithm is presented based off of fully deterministic ordinary differential equations (ODEs). This framework utilizes model predictive control to challenge state of the art correlation scores while retaining the interpretability that comes inherent with a model. Moreover, these computations can be done in real time, thus enabling online experimentation informed by neuronal firing rates. To demonstrate the use cases of this architecture, it is tested on a ground truth dataset courtesy of the *spikefinder* challenge. Finally, the predictive power of the methodology put forward here is underscored by demonstrating its ability to aid in the creation of novel indicators utilized in two-photon imaging.

**Author summary:** We put forward a novel approach to infer when neurons fire as a function of calcium concentration. These calcium recordings are useful for imaging whole populations of neurons, such as those found in the brain, but act only as a proxy for the true underlying spiking occurrences. Moreover, these calcium traces are unfortunately noisy. To uncover the actual firing times we apply modern control architectures to a framework built from chemical reaction equations. The result is accurate, fast enough to provide information in real time, and highly interpretable. More broadly this analysis process aids understanding in contexts where methods of measurement obfuscate the desired ground truth information. Not just capable of inference, prediction is possible thanks to the fully mechanistic formulation provided here. Specifically, we may quantify how indicators, which allow the tracking of calcium, behave in the presence of differing conditions. In more general terms, this framework also enables understanding of the tools used to measure the desired underlying signals.

## Introduction

Information processing within populations of neurons is frequently uncovered via two-photon imaging [1]. This method indirectly tracks concentrations of intracellular calcium ions by their fluorescence, which is a noisy realization of the underlying calcium signal [2]. An action potential within a neuron results in an uptake in intracellular calcium, in turn yielding an increase of fluorescence as calcium ions bind to an indicator. Downstream analyses are frequently concerned with true spiking times, not fluorescent traces. Converting between a fluorescent trace and true spiking times is nontrivial, coming with many computational difficulties. Amongst them are disparities in the temporal resolution of imaging compared to neuronal dynamics, noise in fluorescence, nonlinear relationships between calcium indicators and fluorescence, and biological parameters unknown a priori [3].

This work is concerned with inferring a true firing signal from noisy time series calcium imaging data. State of the art methods approach this in a handful of different ways. Ranging from supervised learning [4], generative methods [5], and nonnegative deconvolutions manifesting as convex optimization problems [6, 7], a litany of theory has arisen as a means to uncover the true spiking occurrences of a neuron. The works of [4, 8] and later [9] present thoughtful surveys of leading methods, while suggesting their preferred approach.

Of particular interest as of late is the ability to analyze systems of neurons in real time. Closed loop investigation of neural circuitry requires quick data processing and carefully crafted experimental conditions. With these conditions met, experiments can deliver sensory stimuli informed by system-wide neural dynamics. For example, a closed loop strategy was employed to tease out how activity in visual processing regions are connected to specific brain states [10]. Recent work from [11] has shown how real time inference of neural activity can inform what stimulation should be supplied to the neurons undergoing imaging, though these authors use calcium traces directly as their measures of activity. Model prediction control (MPC), tracing back to 1960s [12], has caught attention in biological control system, resulting in high levels of control accuracy [13, 14]. In this type of controller, a model, often based on ordinary differential equations, is used to predict optimal control strategy via optimization over a finite time window of data. With only a handful of methods fast enough to be considered real time [7], we aim to add to this body of existing methods by employing tools such as MPC from the controls community.

## Methods and Models

This section utilizes a chemical reaction network to build a model, then discusses how model predictive control (MPC) can be tailored to infer underlying firing rates.

### Chemical Reaction Formulation

Starting with first principles, we examine the chemical reaction network theorized to dictate the interplay between a calcium ion, a calcium indicator, and a bound (or fluoresced) calcium indicator. These quantities will be denoted [Ca^2+^] = *x*(*t*), [CI] = *y*(*t*), and [CI^*∗*^] = *z*(*t*) respectively, where [*·*] denotes concentration. Intracellular calcium binds to an indicator at a rate of *k*_*f*_. Simultaneously these compounds may unbind at a rate of *k*_*r*_, leaving the indicator and calcium ion free within the cell once again. This gives

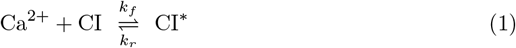

With this chemical reaction network, we may utilize the law of mass action, which posits the rate of a reaction is proportional to the product of reactant concentrations [15, 16].

Supposing calcium indicators do not passively diffuse out of the cell, the total concentration of calcium indicator is time invariant. That is the sum of concentrations of the indicator, both bound and unbound, should remain constant. Call this value *L*. Then for all time *t*

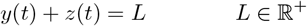

and the equations that follow from (1) may be reduced to

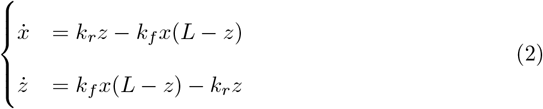

Note these ODEs are nonlinear due to the product of *x* and *z* appearing in both equations. Motivated by the observation that a neuron’s firing results in an uptake of calcium [17], and ultimately is the dominating force controlling the balance of *x* = [Ca^2+^], we add in a continuous function of time *s*(*t*) to represent firing rate at a given time *t*, scaled by some constant *α*. We finally note calcium ion *x* could passively diffuse out of the cell, and subtract a *γx* term to account for this, arriving at the final governing ODE system

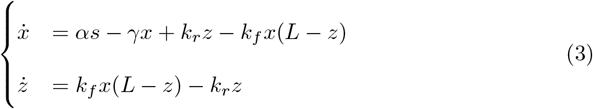

where

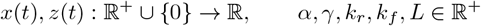

In this phrasing, the question of inferring spiking rates as a function of calcium imaging has been cast as an inverse problem, in which we learn the control signal responsible for driving the dynamics witnessed. A table of parameter values and their biophysical interpretation can be found in the supporting information section, S1 Table.

Justification that this system is stable for all achievable values of *s* can be found in the supporting information section, S1 Appendix.

### Model Predictive Control and the Neuron

MPC is an optimal control technique. This well tested method computes necessary control actions that minimize a cost function and adhere to an underlying model [18], in our case a constrained set of ODEs defined in equation (3). This control action is computed over a finite horizon, enabling changes to actuation that update with real time information. For all simulations in the results section, the open-source MPC package do-mpc [19], implemented in python, was utilized.

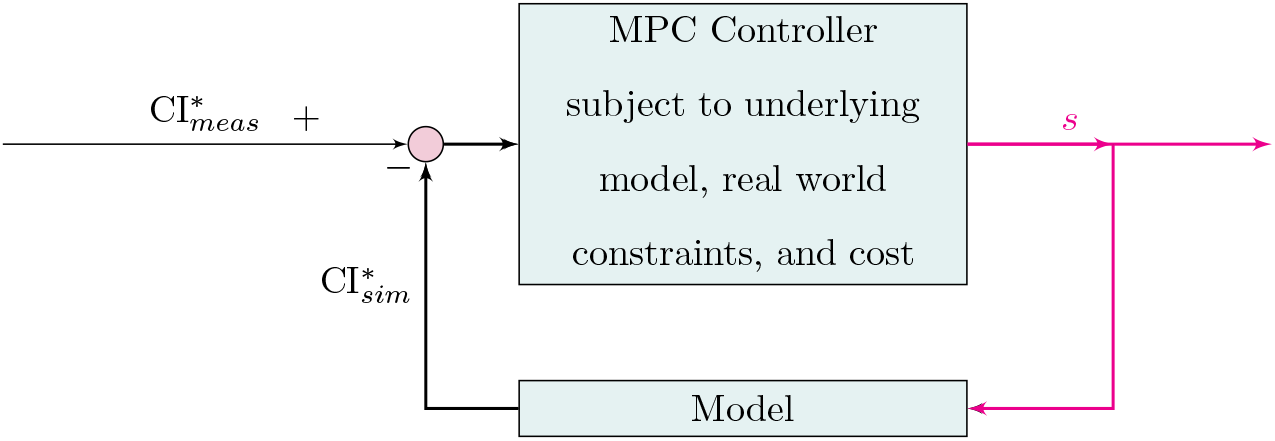

In the above block diagram, 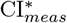 denotes the measured amount of fluoresced indicator, while 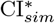 represents the simulated amount of this quantity as dictated by the model (3). Per the definition of MPC, we require a cost defined over our horizon of *n* timesteps. For simulations to follow, we consider the following minimization problem

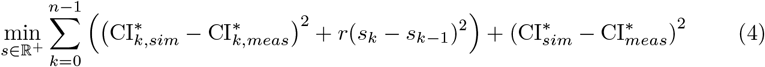

for parameter *r ∈* ℝ^+^. The first squared term enforces our simulated and measured calcium indicators track one another on a finite horizon, and *s*_*k*_ *− s*_*k−*1_ penalizes changes in *s*, essentially limiting the derivative and encouraging smoothness. All simulations to follow take *r* = 0.01 as regularization term. Other authors have had success with the similar cost functions, sometimes using a *l*_0_ or *l*_1_ penalty on *s* instead of the quadratic cost above [6, 7]. Observe that the final term in (4) acts as terminal penalty function, which aids the effectiveness of MPC. An improperly chosen terminal penalty function may degrade performance and potentially destabilize the closed-loop system. Although increasing the prediction horizon can improve performance, this significantly raises computational costs. A well-established solution to these issues involves selecting the terminal penalty function as the infinite-horizon value function that satisfies the dynamic programming equations [20, 21], as is done above. The horizon length *n* = 6 was utilized for all simulations.

In order to partially address the noisy fluorescence signal, a shifted sigmoidal filter

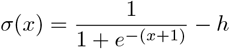

is applied to 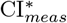. To avoid saturation of this filter, e.g., 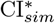 values becoming stuck at 1, a vertical shift downward of *h* = 0.15 is additionally used. If saturation still occurs, a more aggressive vertical shift of *h* = 0.25 is then applied and computations are restarted. The nature of a sigmoidal curve minimizes contributions from extreme values, mapping them closer to 0 or 1, while retaining a linear regime for middle of the pack measurements. More advanced filtering techniques to address stochasticity inherent in the system could undoubtedly improve this methodology, this is discussed briefly in the following section.

## Results and Discussion

With a model formulated and the tools of inference defined, this section benchmarks performance using the *spikefinder* [22] challenge dataset.

### *Spikefinder* Validation

This dataset contains not only time series calcium imaging data, but recorded “ground-truth” spikes via electrophysiology. It is one of the few public datasets to contain simultaneous recordings of spiking times and fluorescent traces, allowing for validation of many of the models discussed in this document. An application of our methodology, henceforth called the MPC approach, to a particular neuron produces a time series of simulated bound calcium indicator 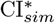 and spiking rates *s* as a function of measured indicator 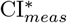, visualized below in Fig 1.

**Fig 1.**
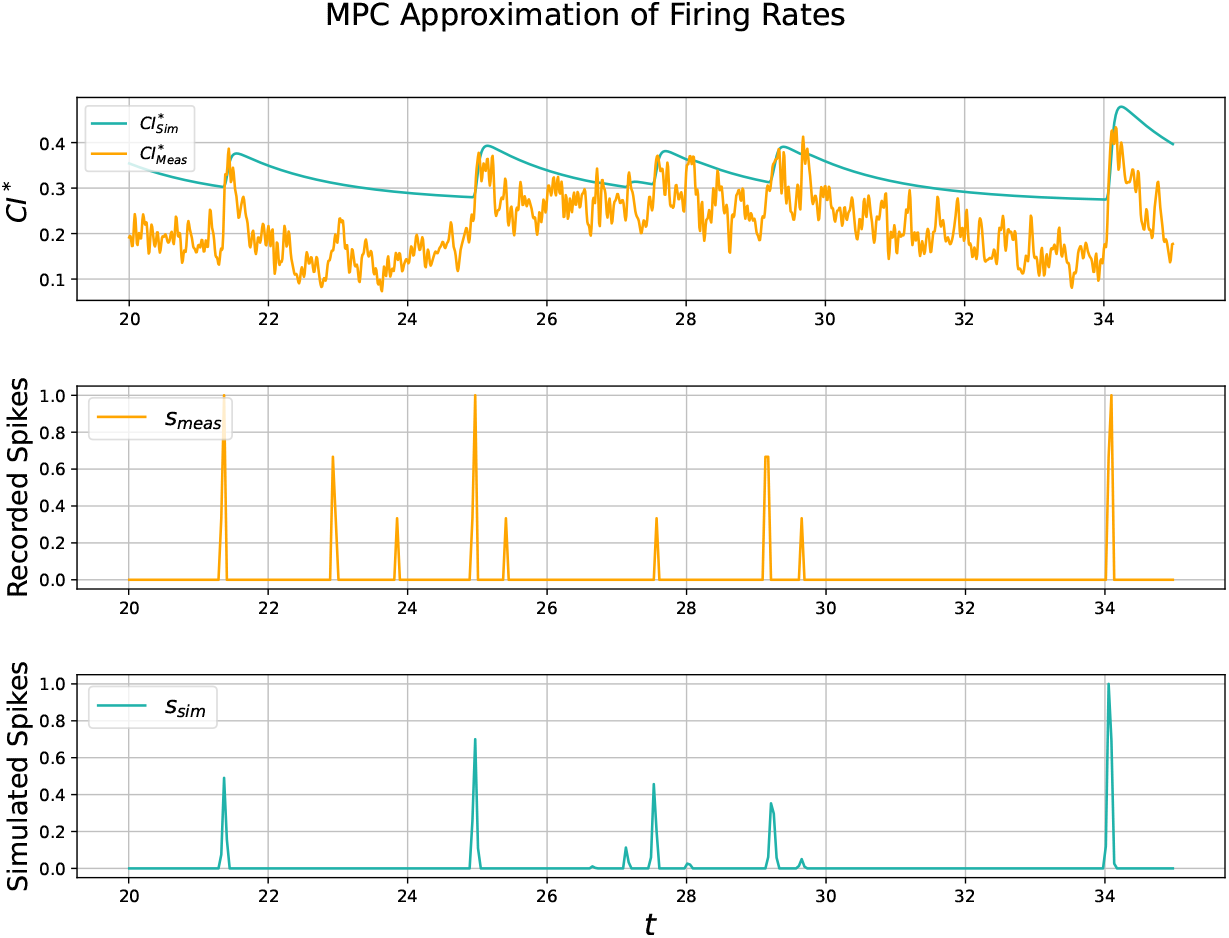
Visualization of Approximations. Pearson correlation coefficient is 0.694 for this particular 15 s subset of test data set 1, neuron 0. The whole 12 min recording scores a 0.491. Spikes normalized to be within [0, 1] for ease of viewing. A few false spikes are present, while others are missed.

The primary metric valued during the *spikefinder* challenge was Pearson correlation coefficient. While emphasis is still placed on correlation, alternative methods of measuring spike synchrony have been put forward such as Victor-Purpura and van Rossum metrics [23–25]. Since spikes are inherently discrete events, for comparison of two spike trains to be computationally tractable and informative we downsample in accordance with the *spikefinder* challenge methodology.

Beginning with correlation coefficient, we compare against the state of the art algorithms STM [4] and Oasis [7] in Fig 2. Here STM serves as a representative of supervised methods, while Oasis is a NND algorithm. Note the work of [7] is another method that is certainly fast enough for real time applications, alongside the MPC approach put forward here. Oasis was eventually benchmarked on the spikefinder dataset by [9], this NND method has a slight advantage in correlation score, but does not have the same amount of interpretability as the MPC approach.

**Fig 2.**
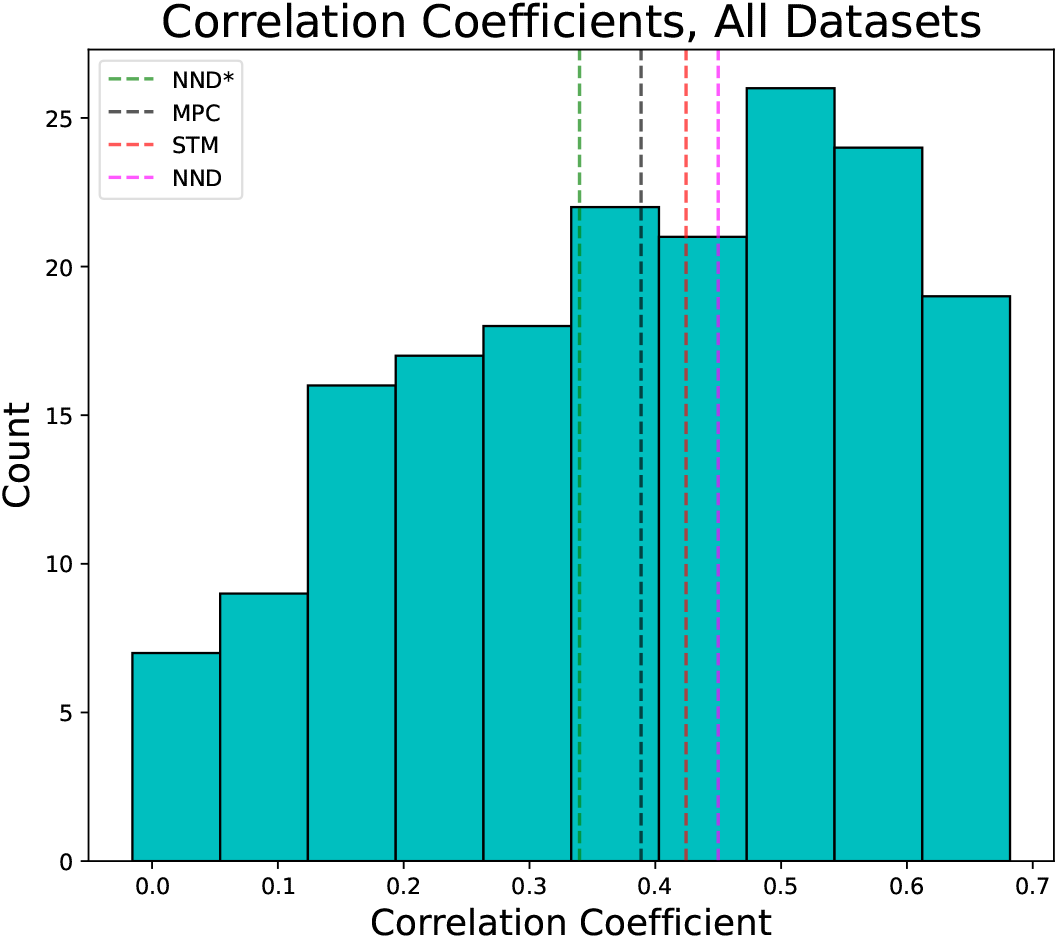
Comparison to state of the art. Histogram of correlation coefficients across all data sets, both train and test. The red dashed line denotes the mean of the STM algorithm, green denotes Oasis supplied with information about the indicator, pink denotes Oasis with no information about the indicator, and the blacked dashed line denotes the mean of the plotted dataset. Means presented are across all data sets, train and test, as well. Resultant firing signals downsampled by a factor of 4 in accordance with the *spikefinder* challenge, yielding a bin width of 40 ms.

As can be seen in Fig 2, our Δ mean relative to STM is only 0.03. While STM does perform better, it requires ample training datasets. The authors of STM note their algorithm does have the capability to perform well when applied to datasets not seen during training, but this is after a wide variety of datasets have already been processed. For the most accurate predictions, such as the results that produced the mean in figure 3, STM is trained on datasets similar to its eventual testing datasets. Put more concisely, STM’s distribution of training data resembles the distribution of testing, at least for the spikefinder challenge. When considering specific datasets the difference in performance is less clear. There exist individual datasets on which the STM algorithm performs notably better than this MPC approach and vis versa. For example, consider Fig 3.

**Fig 3.**
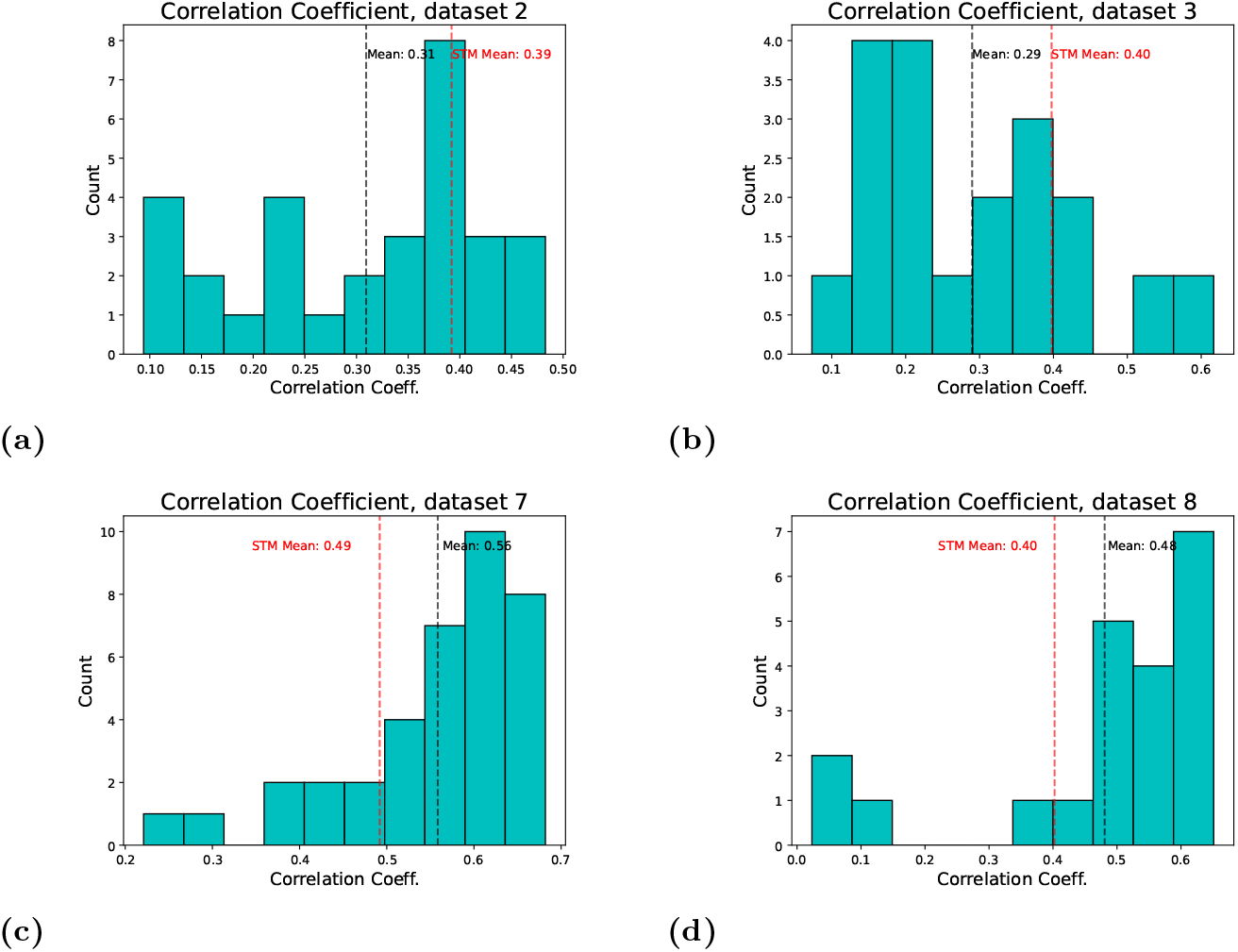
Comparison to STM on specific datasets. Histogram of correlation coefficients computed for four different datasets. (a) and (b) both showcase STM outperforming the MPC approach, while (c) and (d) show the MPC approach outperforming. Datasets 2 and 3 had both train and test data available - both are included. For datasets 7 and 8 only training data was made available.

**Fig 4.**
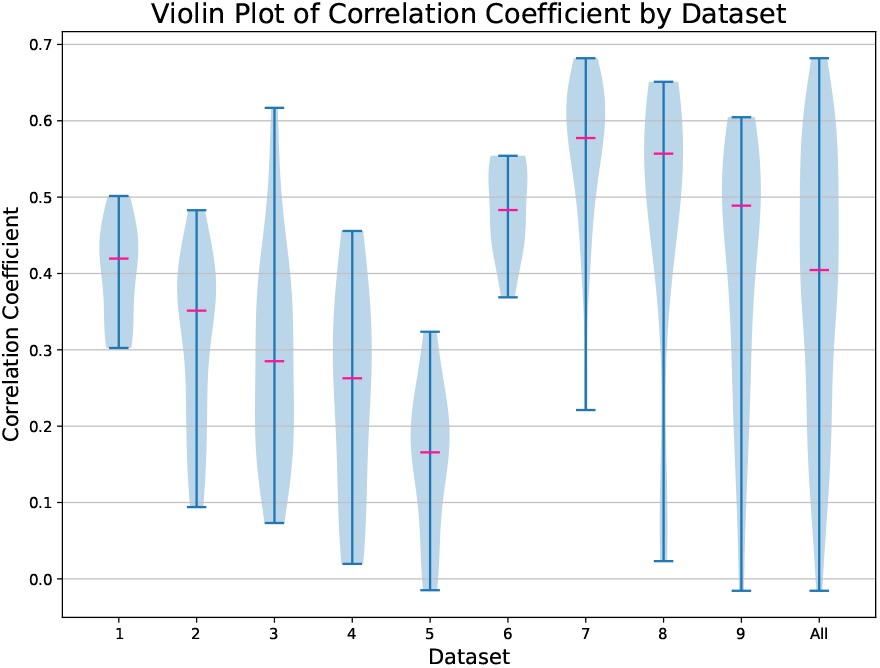
Violin plot of correlation score. Pearson correlation coefficients from MPC approach organized by data set, 40 ms bin sizes. The violin plot uses kernel density estimation to compute an empirical distribution of the samples by dataset. Medians of MPC approach are included in pink.

The MPC algorithm does not require training in the same way a supervised approach such as STM would. Rather, parameters may be selected a priori from known chemical reaction constants, however this undoubtedly results in a sub-optimal parameter regime. For the simulations shown in this section parameters were selected for their biophysical relevance then manually adjusted on training sets in order to maximize correlation coefficients. Thus there is a small training phase, though much shorter and less efficient than that of a neural net’s. With enough biophysical insights, an auto-calibration of parameters should be possible similar to that of MLspike [5], completely alleviating the need to hand train. STM has an auto-calibration functionality, but it requires introducing stimulus and spike history dependence [4].

Prior to examining the aforementioned metric Victor-Purpura distance [23], recall that lower scores are better. In Fig 5 we compare the naive Oasis, meaning Oasis supplied with no information, to the MPC approach. The cost of moving a spike a single timestep is 1 for this simulation.

**Fig 5.**
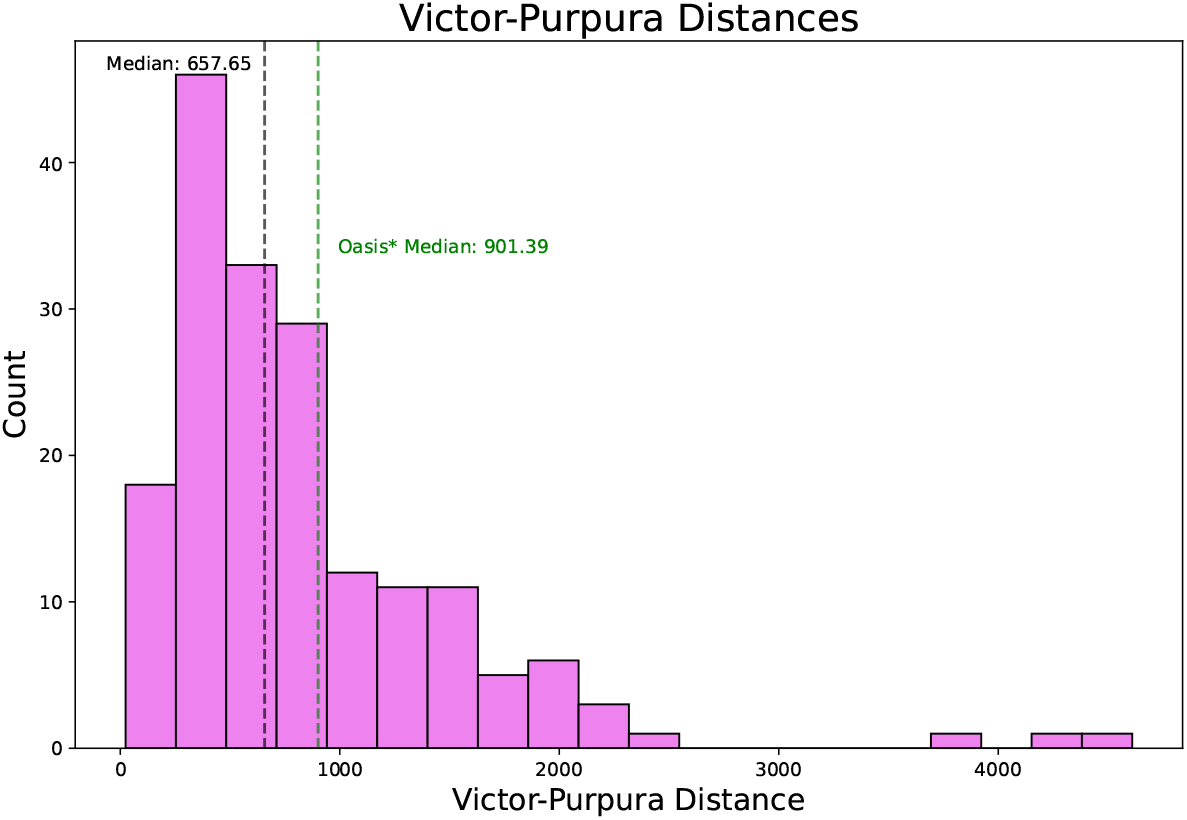
Comparison of Victor-Purpura distance. Victor-Purpura distances across all datasets. Median value of MPC approach shown with a black dashed line, while the median of Oasis (supplied with no information) is shown in green. Since lower is better, we outperform the naive oasis method. A single outlier at 8000 has been excluded from the plot for ease of viewing, leaving 178 different samples visualized in this histogram. Medians were computed with all samples included.

### Reflections on Noise

Though this MPC approach does not require training in a conventional neural net sense, careful selection of parameters is still imperative. This can be seen by considering Figs 6 and 7 in tandem.

**Fig 6.**
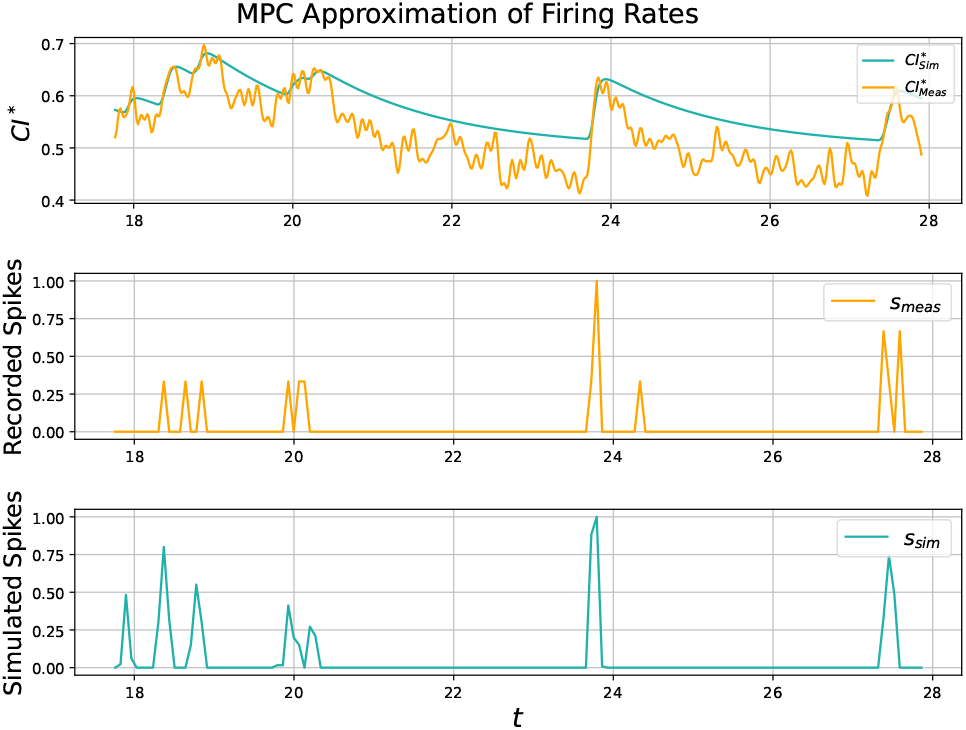
Pearson correlation coefficient for this specific 10 s subset is 0.61, though the whole 12 min recording scores a 0.50. Spikes normalized to be within [0, 1] for ease of viewing. Though most are detected, one is missed at around 24s, with timings slightly off for other spikes.

**Fig 7.**
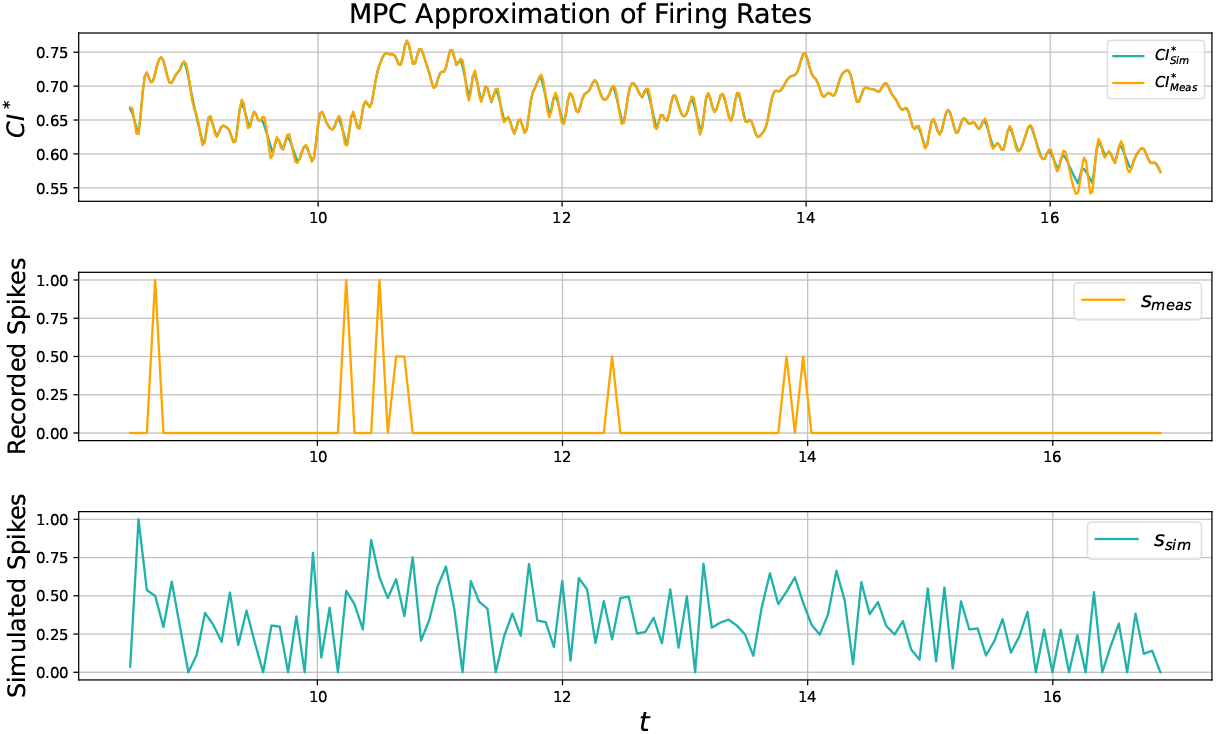
Noise in 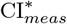 causes the MPC approach to record many false spikes. Correlation coefficient is 0.19

In Fig 6 near optimal parameters are utilized, which incorporates a slow decay rate. This slow decay rate abstracts away noise in the fluorescence signal by avoiding small increases not worthy of a spike. In contrast, Fig 7 has parameters selected to encourage almost perfect tracking of *CI*_*meas*_. While this is easily accomplished via the MPC algorithm, the resultant signal is riddled with false spikes, and thus has a low correlation coefficient. This is because under these circumstances the algorithm tracks the *noise in imaging* instead of the true underlying calcium signal. As mentioned in the introduction, fluorescence is a noisy realization of the true calcium trace, and as such the MPC approach would certainly be improved by incorporating a nonlinear conversion between normalized fluorescence and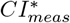. Authors of MLspike [5] incorporate such ideas to great success. In contrast our current formulation equates these two quantities.

## Remarks on Real Time Control

Regarding the usability of methods presented in this document for online control, when training is accounted for only our MPC approach and the NND algorithms are fast enough or structurally able to complete in real time. That is, for all solutions presented the MPC algorithm was able to predict spiking signals in equivalent or less time than the duration of the recording. Though these signals were computed offline, the methodology needs no adaption to run out of the box in real time.

While Jewell’s updated methodology (along with other efficient NND approaches) can impressively compute spiking times for data sets with as many as 100,000 time instants in mere seconds on a laptop, this first requires knowledge of a parameter or training to be accurate. In Jewell’s approach the relevant parameter was discovered by training on the first half of a data set then testing on the second half, suggesting training is still important if high accuracy is desirable. It is worth noting the authors have a way to infer a reasonable parameter regime, though this was not used for testing. In the case of the suite2p implementation of Oasis [9], training is not required but knowledge of the system must be supplied in some way to bring scores in line with means presented in plot 2. Otherwise, the means will be closer to the naive Oasis* instead of the more accurate Oasis. NND models complete on the order of milliseconds however, which is notable considering their stand out accuracy.

The MPC algorithm takes longer than these NND methods; with run times almost always 10-30% less than the duration of their recording. However there is a rich legacy of MPC used for real time control, with many packages optimized for speed and ease of implementation, and as such is relevant in this context for its accuracy and tractable nature. Moreover, this MPC approach allows for other quantities of interest to be inferred, such as the concentration of unbound calcium indicator within the cell. This is a result of our model’s rooting in biophysical mechanisms.

## Biophysical Insights

With the MPC approach and the model underlying it justified, analysis of the system may be considered. Thus far we have been told the status of state variables and inferred *s* from this signal. Reaping the benefits of a fully mechanistic model, we may now suppose *s* and analyze the impacts on state variables.

To accomplish this, we perform frequency response analysis on the system of ODEs (3),

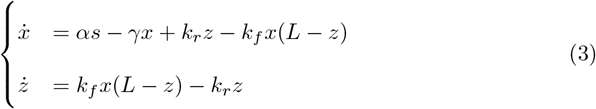

then move to show how this process can be reverse engineered such that an indicator may be designed to work within a specific regime of frequencies.

Since our ODEs are nonlinear many of the elegant results pertaining to frequency response of linear systems is unavailable. Instead, we must suppose a range of frequencies at which *s* oscillates then note the impacts on our state variables of interest.

Specifically, take

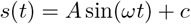

for parameters *A, ω, c ∈* ℝ. This process is showcased for a single *ω* in Fig 8. The response in calcium ion [Ca^2+^] = *x* and indicator [CI^*∗*^] = *z* follow from the supposed *s* in the leftmost plot.

**Fig 8.**
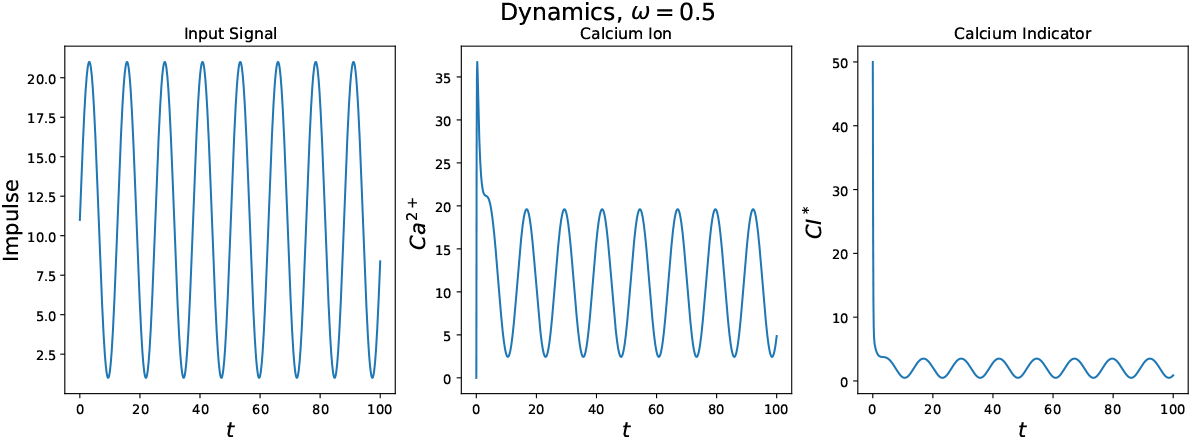
Parameters: *A* = 20, *c* = 1, *ω* = 0.5. Notice after transient dynamics the resultant amplitudes for the calcium ion, indicator are not as large as the amplitude of the *s* signal. This suggests these state variables do not keep up with changes in *s* at this frequency.

To avoid confusion, note *s*(*t*) is the firing rate at given time *t* and is measured in Hz. Here we examine how frequently *s* must change before state variables begin to lag. This is determined by the parameter *ω*, which determines the frequency of the *s* signal, also measured in Hz. For the previous example, sin(0.5*t*) has period 4*π* and thus oscillates at a frequency of 1*/*(4*π*) *≈* 0.080 Hz between the values of 1 and 21 Hz.

To evaluate the ODE system’s performance for a range of *ω*, we perform simulations akin to those done in Fig 8. However after transient dynamics conclude, we note the ratio of amplitudes between *s* and our state variables. This process is showcased in Fig 9. In this way we are able to tease out at which frequencies our state variables are able to reasonably respond to.

**Fig 9.**
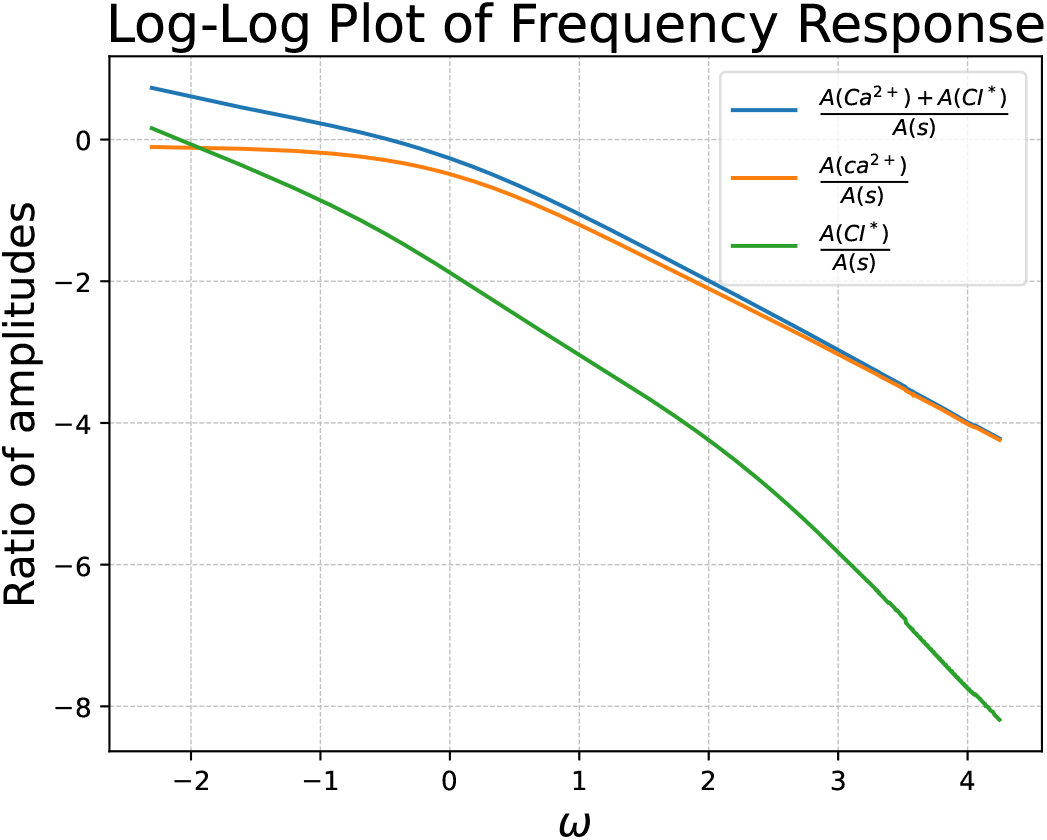
Frequency response analysis of commonly used indicator GCaMP6s, *k*_*f*_ = 0.0514, *k*_*r*_ = 7.6, *L* = 30. Shows for higher frequency oscillations in firing rate calcium dynamics cannot keep up, evident in the amplitude of *s* dominating the amplitude of calcium. Blue shows the total amount of calcium in the system, both bound and unbound. Orange depicts the unbound calcium ions themselves, while green shows the calcium ions bound to an indicator. Justifies utilizing firing rate as the metric for neural activity.

The above calcium indicator keeps up with changes in firing rate poorly for higher frequency oscillations, highlighting the need to convert recorded calcium indicator quantities to firing rate for certain downstream analyses in which a neuron may change the rate at which it fires frequently.

This not only has the power to analyze frequency response for existing indicators, but the potential to aid in the development of novel indicators. For example, if we desired an indicator which operated better for smaller values of *ω* than GCaMP6s, we might try setting new *k*_*f*_ and *k*_*r*_ then examine the resultant frequency response plot. To this end, consider Fig 10, which achieves precisely this by utilizing *k*_*f*_ = 0.01 and *k*_*r*_ = 10 as reaction rates for this toy example.

**Fig 10.**
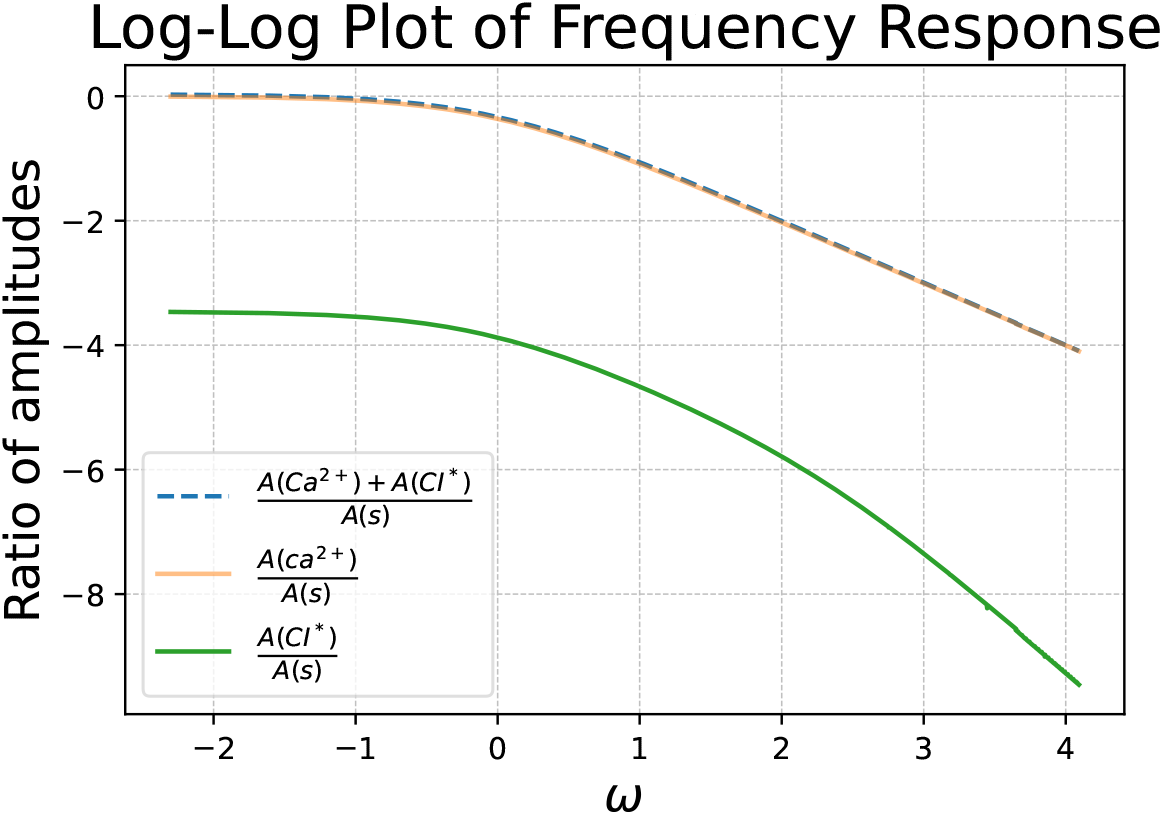
Frequency response of a hypothetical indicator. All parameters are held constant aside from *k*_*f*_ = 0.01 and *k*_*r*_ = 10. Notice an improvement in ratio of amplitudes at small *ω* when compared to figure 9. A drop off still begins at *ω* = 0 or about *e*^0^ = 1 Hz.

The task at hand then becomes creating an indicator with binding affinity *k*_*d*_ = *k*_*f*_ */k*_*r*_ = 0.01*/*10. If additional control over indicator dynamics is desirable, each parameter in (3) may be tuned such that the frequency response is satisfactory.

Constructing an indicator which agrees with all parameters in our governing ODE equations is likely difficult, but nonetheless this has the potential to serve as an approximate guide for the creation of future indicators.

As discussed in [26], general performance criteria to consider in the construction of GECIs include

i. large dynamic range
ii. high calcium sensitivity
iii. faster response kinetics
iv. linear response properties.

The frequency response analysis pipeline outlined in this section could prove invaluable to evaluating the above performance metrics.

## Future Work

An auto-calibration of parameters *α, γ* in equation (3) would increase the usability of this software. To accomplish this, some connection between these parameters and the dynamic qualities of the fluorescent trace must be established. In the style of suite2p, *k*_*f*_ and *k*_*r*_ could be chosen by simply supplying knowledge of the indicator’s speed.

Finally, any of these terms could be learned through supervised learning if labeled data is available.

As another avenue for improvement, consider the terminal penalty function of (4). This function can also be approximated via the Q function in reinforcement learning (RL) [27], presenting an opportunity for future directions of this work in applying RL-based approaches for efficient MPC design. Finally, a more sophisticated treatment of noise would increase accuracy further.

This could be achieved by coupling another ODE connecting fluorescence to calcium, as is done by MLspike, or by employing particle filtering methods. Sequential Monte Carlo methods appear uniquely suited to address the stochasticity present in the system [28].

## Conclusion

In this paper, we propose a novel modeling framework and strategy to infer underlying firing rates of neurons provided calcium imaging traces. This first principles approach underscores the potential of adapting modern control architectures to this space of problems. Accurate enough to challenge state of the art methods, this formulation joins the select few methodologies capable of providing real time spiking information. This fully mechanistic model is highly interpretable and capable of not just inference, but can uncover and predict biophyiscal nuances as they relate to hyperparameters such as reaction rates of indicators.

## Supporting information

### S1 Appendix. Stability analysis

In this appendix the governing ODE system (3) is shown to be stable for all realistic values of *s*. Begin the stability analysis by considering the intersection of nullclines. Let 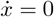, *ż* = 0. After a bit of algebra the coordinates of fixed point (*x*^*∗*^, *z*^*∗*^) is found to be

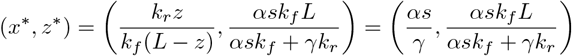

The Jacobian of the system is given by

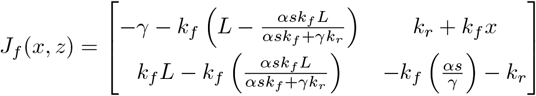

Examining the eigenvalues of this matrix upon evaluation at (*x*^*∗*^, *z*^*∗*^), the condition put on s in order yield two negative eigenvalues, and hence stability, is simply that *s ∈* ℝ. Note these bounds are found from numerical approximations, as an exact solution becomes intractable.

Since the range of *s* is restricted to the positive reals, it follows that we have stability for all attainable values of *s*. To visualize that these ODEs produce a stable fixed point as a function of *s* consider the following phase diagram, in which *s* is fixed to be 30. Since our fixed point is a function of *s*, it follows that taking *s* as a time varying function *s*(*t*) would simply shift the location of our attractor. Taking parameters *α* = *γ* = 1, *k*_*r*_ = 7.6, *k*_*f*_ = 0.05135, *L* = 100, *s* = 30, we would expect our fixed point to be at

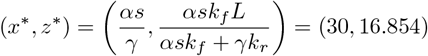

This is visualized in Fig 11.

**Fig 11.**
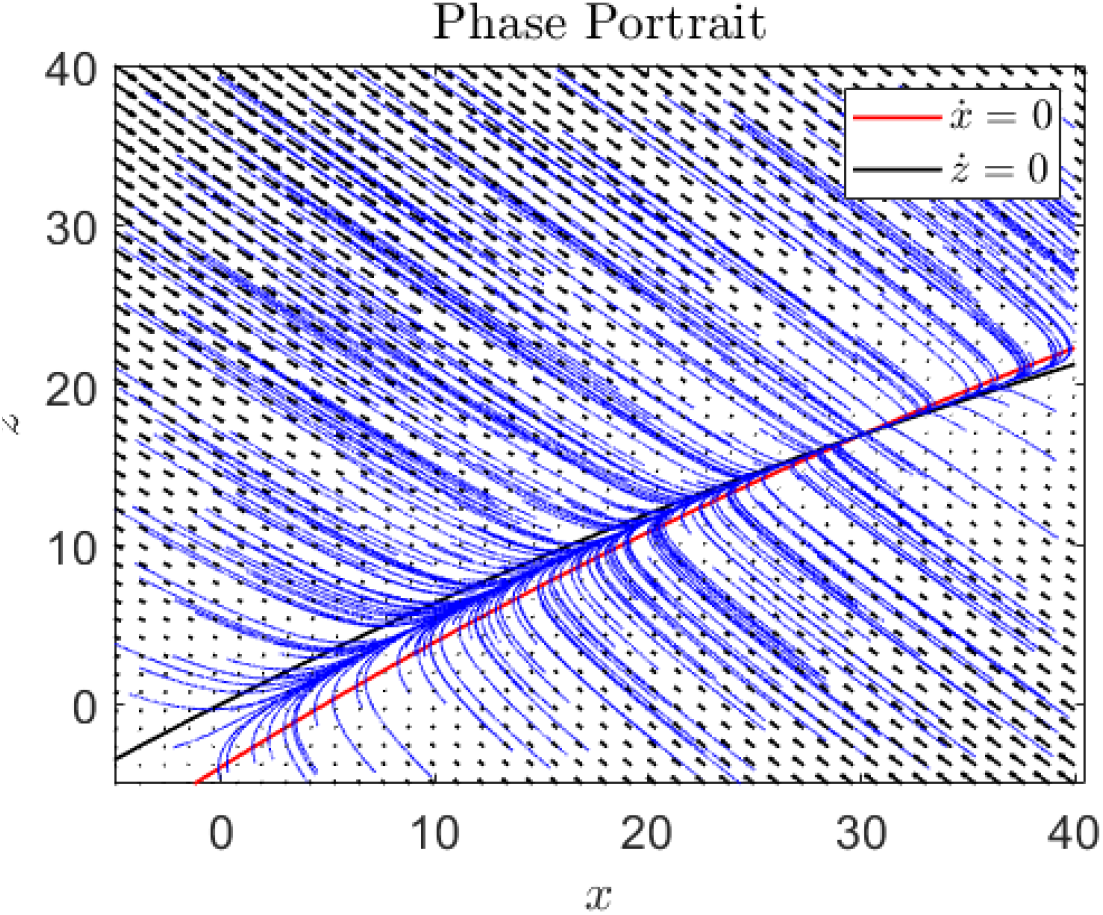
Flows of equation (3). For this simulation, *α* = *γ* = 1, *k*_*r*_ = 7.6, *k*_*f*_ = 0.05135, *L* = 100, *s* = 30. Note our fixed point is, as was expected, at (30, 16.854).

## S1 Table Parameter values and physical interpretation

The following tables provide physical interpretation and units for governing equation (3). Values provided were utilized the in *spikefinder* simulations.

**Table 1.** Datasets 1-3, and 6-9.

| Parameter | Value | Units | Description |
| --- | --- | --- | --- |
| $k_r$ | 0.1 | $\mu M^{-1} s^{-1}$ | Forward reaction rate |
| $k_f$ | 10 | $s^{-1}$ | Reverse reaction rate |
| $\alpha$ | 16.66 | $\mu M/\text{spike}$ | Weighting of firing |
| $\gamma$ | 1.366 | $s^{-1}$ | Passive diffusion scalar |
| $L$ | 100 | $\mu M$ | Total indicator available |
| $\Delta t$ | 0.01 | $s$ | Simulation step size |

**Table 2.** Dataset 4.

| Parameter | Value | Units | Description |
| --- | --- | --- | --- |
| $k_r$ | 0.1 | $\mu M^{-1} s^{-1}$ | Forward reaction rate |
| $k_f$ | 10 | $s^{-1}$ | Reverse reaction rate |
| $\alpha$ | 10 | $\mu M/\text{spike}$ | Weighting of firing |
| $\gamma$ | 0.73 | $s^{-1}$ | Passive diffusion scalar |
| $L$ | 100 | $\mu M$ | Total indicator available |
| $\Delta t$ | 0.01 | $s$ | Simulation step size |

**Table 3.** Dataset 5.

| Parameter | Value | Units | Description |
| --- | --- | --- | --- |
| $k_r$ | 0.2 | $\mu M^{-1} s^{-1}$ | Forward reaction rate |
| $k_f$ | 10 | $s^{-1}$ | Reverse reaction rate |
| $\alpha$ | 10 | $\mu M/\text{spike}$ | Weighting of firing |
| $\gamma$ | 0.73 | $s^{-1}$ | Passive diffusion scalar |
| $L$ | 100 | $\mu M$ | Total indicator available |
| $\Delta t$ | 0.01 | $s$ | Simulation step size |

## Acknowledgments

Thank you to the Gomez lab group, in particular Ksenia Zlobina, for insightful discussions pertaining to modeling and analysis.

## Notes

### Competing Interest Statement

The authors have declared no competing interest.

### Summary of Updates

Order of authors erroneously differed from the actual manuscript, this is now rectified. All author's Orcid IDs are now linked as well.

